# Soft silicone surface stiffening by oxidation upon deep UV treatment as characterized using nanoindentation

**DOI:** 10.64898/2026.06.19.733410

**Authors:** Amanda Wilder, Zaria Booth, Caden Obermeyer, Saika Sharmin, Venkat Maruthamuthu

## Abstract

Silicones are elastomers that have a wide variety of uses, including biomedical applications such as the coating of biomedical devices and as implants. Soft silicones with mechanical properties similar to those of biological tissues have particularly gained use as substrates for cell culture in mechanobiology studies. In this context, it would be desirable to be able to alter their surface mechanical properties with a relatively simple physical treatment. While deep ultraviolet (deep UV) or ultraviolet C (UV-C) treatment has been previously used as a surface treatment method for stiffer silicones formulations, the effect of this treatment on soft silicones relevant for mechanobiology applications is still uncharacterized. We first used nanoindentation to determine the Young’s modulus of two types of soft silicones, Qgel and GEL-8100/Syl (GEL-8100 with Sylgard-184 crosslinker), both with initial moduli in the kilopascal range. We show that nanoindentation in the presence of 1% sodium dodecyl sulfate avoids adhesion between the nanoindentation glass probe and the soft silicones. After deep UV exposure in the presence of air, nanoindentation revealed that the apparent Young’s moduli of the soft silicones Qgel and GEL-8100/Syl increased by 70% and 33%, respectively. The bulk rheology of the soft silicones were not affected, suggesting that this corresponds to a surface stiffening effect with a topical stiffening of at least several hundred kilopascals. Energy-dispersive X-ray spectroscopy results show an increase in the mole fraction of oxygen, consistent with oxidation of the surface. Attenuated Total Reflectance Fourier-Transform Infrared spectra show evidence of Si-OH group formation in GEL-8100/Syl and silicon sub-oxide formation in Qgel. Consistent with this, water contact angle measurements show enhanced hydrophilicity after deep UV treatment. Our results have implications for using soft silicones as substrates in mechanobiology studies and in processes where deep UV light is used in the surface treatment of soft silicones.

## Introduction

Silicones are a broad class of polymers with a -(-O-R_2_Si-)- repeating unit. The presence of the -Si-O-Si-O- inorganic backbone and the -R organic groups bestows them with unique physicochemical properties, as has been widely noted (1, 2). In particular, silicone rubbers are elastomers with a variety of applications (2, 3), including as biological scaffolds and as coatings in biomedical devices (4). Polydimethylsiloxane (PDMS) is perhaps the most notable silicone, and has many important biomedical applications including as implants (5). The most commonly used version of PDMS is Sylgard 184, which is typically used at a composition that yields an elastomer of ∼MPa elastic modulus (6). The high elastic modulus of this version of PDMS, as well as its biocompatibility, have made it ideal for applications such as microfluidics (7–9) and as spatial barriers to restrict or pattern cell culture (10, 11). Using Sylgard 184, PDMS as soft as several kPa can be achieved by using a smaller cross-linker fraction (12), but most studies using Sylgard 184 use it at a composition that yields ‘stiff silicone’ of MPa elastic modulus.

In contrast to this, ‘soft silicones’ are silicone elastomers (typically these are also chemically PDMS or close variants, but with different polymeric chain arrangements) that can readily be prepared with elastic moduli in the sub-kPa to tens of kPa range (13) that is most relevant for mimicking biological tissues, cells (14) or the ECM. Biological tissues, largely comprised of living cells and the extra-cellular matrix (ECM), are viscoelastic materials (15, 16), with elastic moduli in the sub-kPa to the supra-MPa range, depending on tissue type (17). Importantly, the mechanical properties of the ECM critically determine cell behavior, influencing cell survival, migration and differentiation (18). Biomaterials that can mimic properties of the ECM have enabled the engineering of cell responses, guiding of cell differentiation and tissue repair (19). Hydrogels like polyacrylamide gels have also been used as substrates for cell culture (20–23) and have helped decipher cell responses to ECM-like substrate stiffness. However, hydrogels suffer from drawbacks such as drying out in the absence of aqueous media and inducing hydraulic cracks in tissues in applications requiring external stretch (24). Thus, ECM-coated soft silicones have emerged as a useful ECM-mimicking substrate for cell culture in mechanobiology studies (14, 25–28), in a manner complementary to hydrogels. Soft silicones, much like polyacrylamide gels, have mechanical properties that are dependent on their composition, which in turn involves different extents of cross-linking of their constituent polymer chains during curing. However, it is desirable to have a method to alter their mechanical properties after curing, for a given chemical composition. This can enable paired experiments and comparison of the effect of just the altered mechanical property of the substrate, independent of sample-to-sample variations in silicone mixture preparation.

Prior work with stiff silicones suggest that exposure to lower wavelength UV light, especially in the presence of oxygen, can lead to cross-linking of siloxane chains (29) and an increase the hardness of silicone (30–34). Some of these studies considered PDMS based on Sylgard 184 (32, 34), while other studies considered high temperature vulcanized rubber (29) or methyl vinyl silicone (30). The length of UV exposure considered also varied from tens of minutes (acute UV treatment) to thousands of hours (long-term UV exposure). UV light can be classified into UV-A, UV-B and UV-C, with UV-C or deep UV light being of highest energy and corresponding to wavelengths under 280 nm. Deep UV (UV-C) light emitted from a mercury quartz lamp has 185 and 254 nm fractions which, in the presence of oxygen, generate ozone. Thus, deep UV light in the presence of air is similar to an ultraviolet ozone (UVO) treatment. UV-C light has been used for disinfecting biomedical silicone used in facial prostheses (35). It has been previously shown that deep UV treatment of stiff silicones like Sylgard 184 PDMS leads to initial polymer chain scission and a subsequent production of a tens of nm thick silica layer (31–34). One study in particular (31) noted a pronounced increase in oxidation of PDMS after 20 min of deep UV exposure. However, it is unclear if deep UV treatment of soft silicones, which possess a chain arrangement necessarily distinct from stiff silicones, also stiffen upon deep UV exposure in the practical treatment timescales of tens of minutes, since the surface stiffening effect depends on the balance of chain scission and oxidation (32, 34). This is pertinent to investigate, given the increasingly common use of soft silicones as cell culture substrates wherein the mechanical properties of the substrate are a key relevant variable. Change of substrate stiffness with a physical treatment method can facilitate cell culture studies with paired substrates as well as spatial patterning of substrate stiffness.

The mechanical properties of silicone gels can be measured in many ways – from bulk rheology of the silicone that can be characterized with a rheometer (28) to surface measurements with indentors (36). Atomic force microscopy of the surface was used to characterize the relative increase in elastic modulus of a stiff silicone previously (34). While bulk rheology to atomic force microscopy spans a large range of length scales, mechanobiology applications using soft silicones with cells typically involves sensing of the elastic modulus at the micron to tens of microns scale. Thus, in this study, we chose to use a nanoindentor with a probe size of a few microns to characterize the elastic properties of soft silicones before and after deep UV exposure. We consider two types of soft silicone, both based on PDMS, and first show that the large adhesion between the nanoindentor glass probes and soft silicone surface can be eliminated by surfactant presence in the medium. We then show that deep UV exposure in the presence of air can be used to increase the apparent Young’s moduli of both types of soft silicone by tens of percent. We use bulk rheology to show that the stiffening is limited to the surface and estimate the extent of surface stiffening. Finally, we use energy-dispersive X-ray spectroscopy (EDS) to show an increase in the mole fraction of oxygen, consistent with surface oxidation upon deep UV exposure that underlies soft silicone stiffening.

## Materials and Methods

### Soft Silicones

Soft silicones used were based on GEL-8100 (NuSil Silicone Technologies, Carpinteria, CA, USA) and Qgel 300 (CHT USA Inc., Richmond, VA). GEL-8100 was prepared by mixing its A and B components at a 1:1 ratio and including 0.5% of the crosslinking component of Sylgard 184 (Dow, Midland, MI, USA) (28). For the Qgel 300 soft silicone, 1 g of Qgel A was mixed with 2.2 g of Qgel B (i.e., 1:2.2 A:B) (27). For the GEL-8100 soft silicone, 4.5 g of GEL-8100 A, 4.5 g of GEL-8100 B and 1g of Sylgard 184 crosslinker were first mixed to yield a 1:1 A:B ratio with 10% Sylgard 184 crosslinker, and then serially diluted with a 1:1 A:B mixture of GEL-8100 to yield a final composition of 1:1 A:B GEL-8100 with 0.5% Sylgard 184 crosslinker. For both soft silicones, the constituent components were mixed in an aluminum weighing boat using a wooden stick for a period of 2-3 minutes. Then, 4-5 drops (formed by gravity off the tip of the wooden stick) of the prepared mixtures were added and spread on top of a 22 mm x 22 mm glass coverslip and then degassed for 5 min under vacuum. The coverslips with GEL-8100/Syl were placed on a glass slide and cured in an oven at 70 °C for 2.5 hrs. The coverslips with Qgel were placed on a glass slide and cured on a hot plate wrapped in aluminum foil at 100 °C for an hour. Each of the two types of soft silicones used (Qgel and GEL-8100/Syl) were made as 7 independent preparations, with each preparation giving rise to one or more soft silicone samples that were untreated and one or more soft silicone samples that were exposed to deep UV light. The variability from one preparation to another is captured in figure 4, where the mean of the Young’s modulus from each preparation is shown as a data point beside the box plot.

### Deep UV treatment

Samples were exposed to deep UV light in a Novascan UV/Ozone system with a 4” (0.1 m) mercury vapor lamp and equipped with a rapid ozone evacuator/neutralizer at a distance of 10 cm (0.1 m) from the lamp for 30 min. The deep UV intensity was deduced to be 2 mW/cm^2^ (20 W/m^2^) at this distance based on the specifications of the manufacturer (Novascan, Boone, IA). A deep UV treatment time of 30 min was chosen based on prior works (31, 32, 34) that also used deep UV treatment in the presence of air.

### Nanoindentation

Nanoindentation was performed using a Chiaro nanoindenter (Optics11 Life, Amsterdam, Netherlands) mounted on a DMi8 inverted fluorescence microscope (Leica Microsystems, Buffalo Grove, IL). The Chiaro nanoindentor head was equipped with a coarse stage with a 12 mm range and a fine stage with a 100 μm range. The nanoindentation probe consisted of a glass ferrule, an optical fiber and a gold coated cantilever with with a spherical glass tip of radius 26.5 μm and pre-calibrated probe stiffness of 0.47 N/m. Cantilever tip displacement (measured using optical interferometry) and piezo movement were used to obtain the indentation as piezo movement minus the cantilever tip displacement. The load is the cantilever tip displacement times the probe stiffness.

Nanoindentation measurements were taken in phosphate-buffered saline (PBS) or in 1% (w/w) sodium dodecyl sulfate (SDS) in PBS. Each soft silicone sample was indented 6 times, 1000 μm apart along x/y, leading to at least 6 indentations from each independent preparation for the untreated case and similar for the deep UV exposed case. For the matrix scans, soft silicone surfaces were probed using the nanoindentor at points that formed a 5×5 grid, with a grid spacing of 1000 μm. The probe size was chosen to be of the order of 10 μm since that is the scale most relevant for cell-level mechanobiology applications. The cantilever downward speed was chosen to be 10 μm/s, since we found that it was the lowest speed at which the noise in the measurements were not pronounced. Use of 1% SDS in the medium nearly eliminated adhesion between the probe and the silicone as detailed in the Results and Discussion section.

The principles of design of experiments utilized were randomization, replication and blocking: The samples in a given batch that were subject to deep UV treatment were randomly chosen. The measurements were replicated by preparing multiple batches that also allowed us to capture batch to batch variation. Finally, the effect of batch to batch variability on deep UV treatment was blocked by considering paired samples – one untreated and one deep UV treated from each batch.

### Hertz model

The Young’s modulus was obtained by fitting the initial part (up to 1-2 μm) of the loading curve with the Hertz model (37) of indentation:

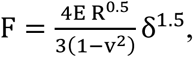 where F is the loading force, δ is the indentation, R is the radius of the indentor, E is the Young’s modulus and ν is the Poisson ratio. Furthermore, the contact radius is given by 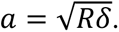 For our measurements *a*,∼5-7 µm. It should be noted that the indentation depth used for the Hertz fit was <10% of the indentor radius as well as <2% of the soft silicone thickness.

### Bulk rheology

Bulk shear rheology of the soft silicones was performed using an MCR-302 rheometer (Anton Paar, Ashland, VA) with an air bearing. The soft silicones were loaded between 25 mm diameter parallel plates for the measurement of storage (G’) and loss (G”) shear moduli. The moduli were obtained as a function of angular frequency in the 0.1–10 rad/s range for 1% strain.

### Scanning Electron Microscopy, Energy-dispersive X-ray spectroscopy and Attenuated Total Reflectance Fourier Transform Infrared Spectroscopy

Scanning electron microscopy (SEM) was performed with a Jeol JSM-6060LV scanning electron microscope, with an accelerating voltage of 15 kV. For this, the soft silicone samples were first coated with a thin layer of a gold−palladium alloy using a Polaron E5100 Series II sputter coater under an atmosphere of argon at less than 0.1 mbar. SEM images were acquired at 30,000x magnification. Energy-dispersive X-ray spectroscopy (EDS) (38) was performed by collecting the X-rays emitted from ∼ 1 μm of the sample top layer in response to the incident electron ray using a Thermo Scientific UltraDry Premium EDS Detector. In EDS, the emitted X-ray energy level in the spectrum is used to identify elements and relative peak intensities of the emitted X-rays is used to obtain element abundance. Attenuated Total Reflectance Fourier Transform Infrared (ATR-FTIR) spectroscopy was performed on both types of soft silicone surfaces (untreated or after deep UV treatment) using a Bruker Alpha II Platinum-ATR device. The soft silicones supported by a glass coverslip base were inverted onto the sample window such that the soft silicones adhered and established good contact enabling immediate signal acquisition. Since the ATR-FTIR signal strength can vary due to the variations in the extent of sample contact, we first normalized the absorbance by scaling to a well-known peak in the bulk that is not expected to vary. Since the ATR-FTIR probes to 1-2 µm depth and the oxidized layer is expected to be an order of magnitude shallower, the peak at 790-800 cm^-1^ corresponding to Si–CH_3_ rocking and Si–C stretching in Si-CH_3_ is expected to be fairly constant in the bulk and so was chosen for normalization. Note that this low wavenumber corresponds to larger penetration depth and hence the signal is expected to be dominated by the bulk, rather than the surface oxidation layer.

### Contact angle measurement

50 µL of deionized water was gently pipetted onto either kind of soft silicone, without or after deep UV treatment that was in turn placed on a horizontal platform. The soft silicones themselves were ∼150 µm thick and were supported by a glass coverslip base. Digital images of the profile view of the water droplet was recorded and analyzed using NIH ImageJ.

### Statistical Analysis

For statistical analysis, a paired t-test was used to compare the Young’s moduli of Qgel without or after deep UV exposure as well as those of GEL-8100/Syl without or after deep UV exposure, with ** indicating p < 0.01 and *** indicating p < 0.001. An unpaired t-test was used to compare the water contact angles on soft silicones without or after deep UV treatment. A paired t-test was considered suitable after a Shapiro-Wilk test for normality yielded p values in the 0.30-0.92 range (depending on the data set) meaning that the null hypothesis that the data are from a normal distribution could not be rejected.

## Results and Discussion

Out of soft silicones that have found use as soft substrates in mechanobiology studies recently (25, 26, 28, 36, 39, 40), we chose two kinds of soft silicones for our work here: Qgel (Qgel made with the ratio of A and B components being 1:2.2) (27, 40) and GEL-8100/Syl (28) (GEL-8100 made with the ratio of A and B components being 1:1 with 0.5% of the crosslinking component of Sylgard 184). When we first attempted to use the nanoindentor (please see Materials and Methods) with a glass spherical tip to indent either sample with PBS as the medium, we found that there was a very high amount of adhesion between the surface of the silicone and the tip of the probe (Fig 1). This not only lengthened the time duration for each indentation cycle, but also precludes use of a simpler indentation model like the Hertz model that assumes negligible adhesion between the probe and the substrate.

**Figure 1.**
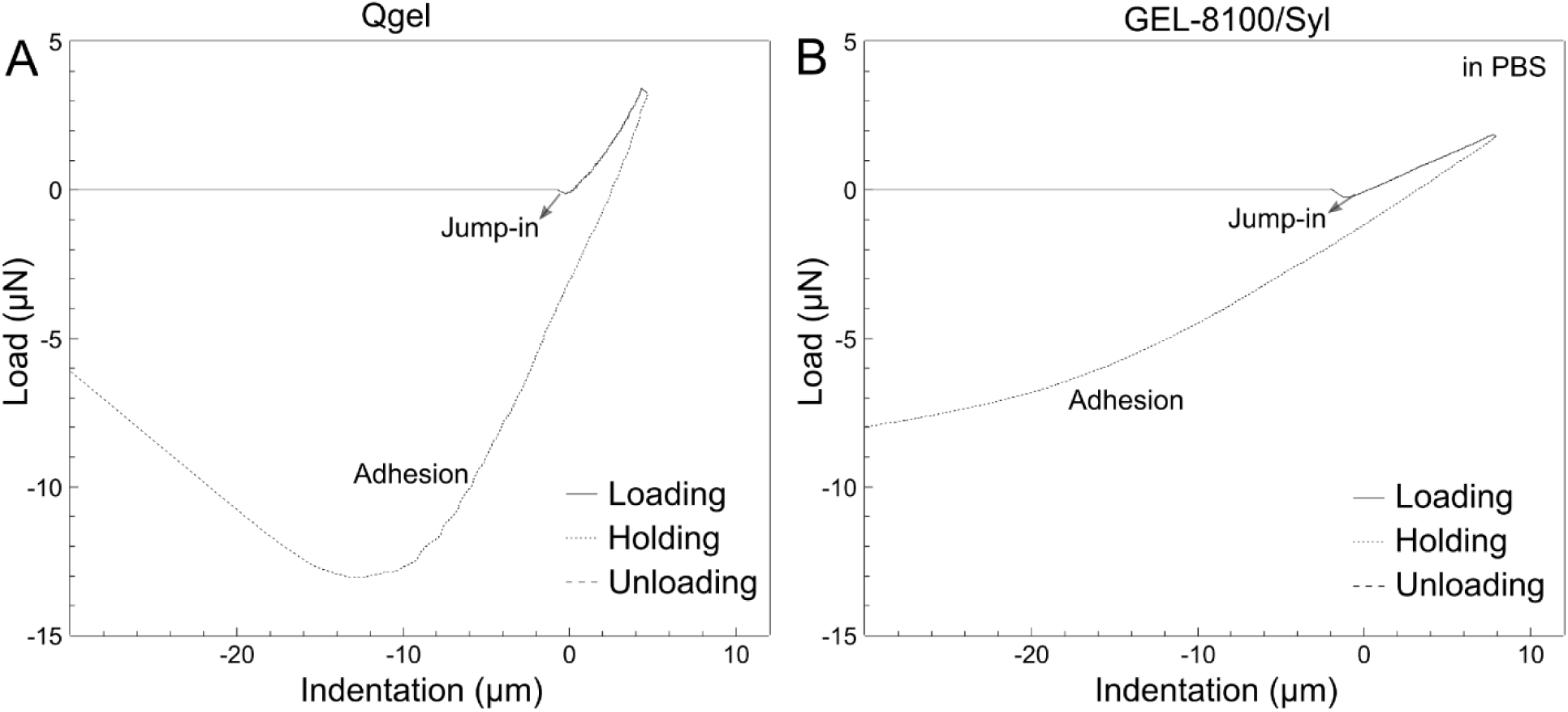
Nanoindentation of both kinds of soft silicone reveals glass probe-silicone adhesion. Load-indentation curves for the indentation of two kinds of soft silicone measured in phosphate-buffered saline (PBS): (A) 1:2.2 Qgel and (B) 1:1 GEL-8100/0.5% Sylgard crosslinking agent. Loading curves are shown as solid lines and the unloading curves as dashed lines. Note the jump into contact as well as the large adhesion seen in the unloading curves.

Based on a previous report (41) that employed a different kind of soft silicone, we hypothesized that using a surfactant such as sodium dodecyl sulfate (SDS) would reduce the adhesion between our soft silicones and glass probes as well. Therefore, we used 1% SDS in PBS as our medium and performed indentations again. We found that the addition of 1% SDS was able to eliminate almost all of the adhesion (Fig 2) for both kinds of soft silicone. We then proceeded to fit the indentation curve (initial 1 μm of indentation for the Qgel and 2 μm of indentation for GEL-8100/Syl) for the loading portion with the Hertz model for elastic materials (Fig 2). The Young’s modulus for Qgel was 20.5 ± 7.5 kPa and the Young’s modulus for GEL-8100/Syl was 6.5 ± 1.5 kPa. It is worth noting that it was previously shown that the Young’s modulus obtained using a Hertz model after the elimination of adhesion agreed with that obtained using the Johnson–Kendall–Roberts model in the presence of adhesion (41). We then wanted to test if deep UV exposure can be used to stiffen the soft silicones.

**Figure 2.**
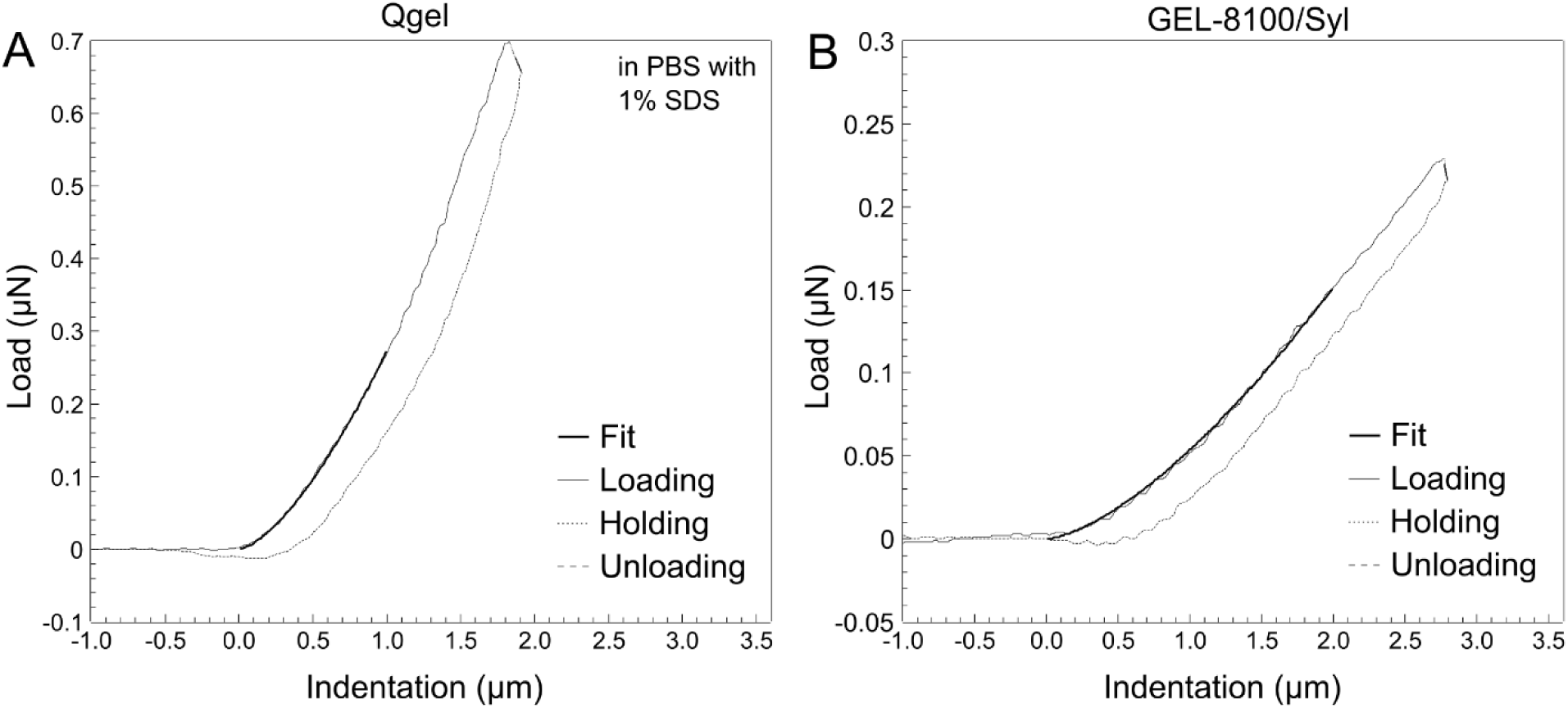
Nanoindentation of both kinds of soft silicone in the presence of surfactant shows elimination of glass probe-silicone adhesion. Load-indentation curves for the indentation of two kinds of soft silicone measured in 1% (w/w) sodium dodecyl sulfate (SDS) in PBS: (A) 1:2.2 Qgel and (B) 1:1 GEL-8100/0.5% Sylgard crosslinking agent. Loading curves are shown as solid lines, unloading curves as dashed lines and the Hertz model fit as bold solid line. Note the near-elimination of adhesion (compared to Fig 1) as seen in the unloading curves.

We exposed both kinds of soft silicones to 30 min of deep UV in the presence of air and then used nanoindentation to determine their Young’s moduli (Fig 3). It is evident from Fig 3A,B that the slope of the load indentation curves increases after deep UV exposure (compared to Fig 2A,B), and that deep UV exposure has a much greater effect on the Qgel than the GEL-8100/Syl. We found that the Young’s modulus for Qgel after deep UV exposure was 34.9 ± 8.7 kPa (Fig 3C) and for GEL-8100/Syl after deep UV exposure was 8.7 ± 2.6 kPa (Fig 3D). This revealed a ∼70% increase in stiffness for Qgel and a ∼33% increase in stiffness for GEL-8100/Syl upon deep UV exposure. Note that, as shown in Fig 3C,D, despite heterogeneity in soft silicone stiffness among independent preparations, each sample preparation (be it Qgel or GEL-8100/Syl) stiffened upon UV exposure. In order to assess the spatial variation of the Young’s modulus along the surface, we performed nanoindentation at points that formed a grid on the silicone surface. As shown in Fig 4 for both Qgel (Fig 4A) and GEL-8100/Syl (Fig 4B), the increase in Young’s moduli upon deep UV exposure is greater than the undulations in Young’s moduli along the surface due to point to point variation. In order to assess if the spatial variation in Young’s moduli observed in Fig.4 are due to spatial heterogeneities or experimental noise, we prepared a Qgel sample and measured the Young’s modulus from 11 indentations at the same location. This yielded a mean value of 10.0 kPa and a standard deviation of 0.5 kPa, i.e., a coefficient of variation of 5%. This suggests that the spatial variation in Young’s moduli within each of the scans in Fig. 4 is likely due to spatial heterogeneity rather than experimental noise.

**Figure 3.**
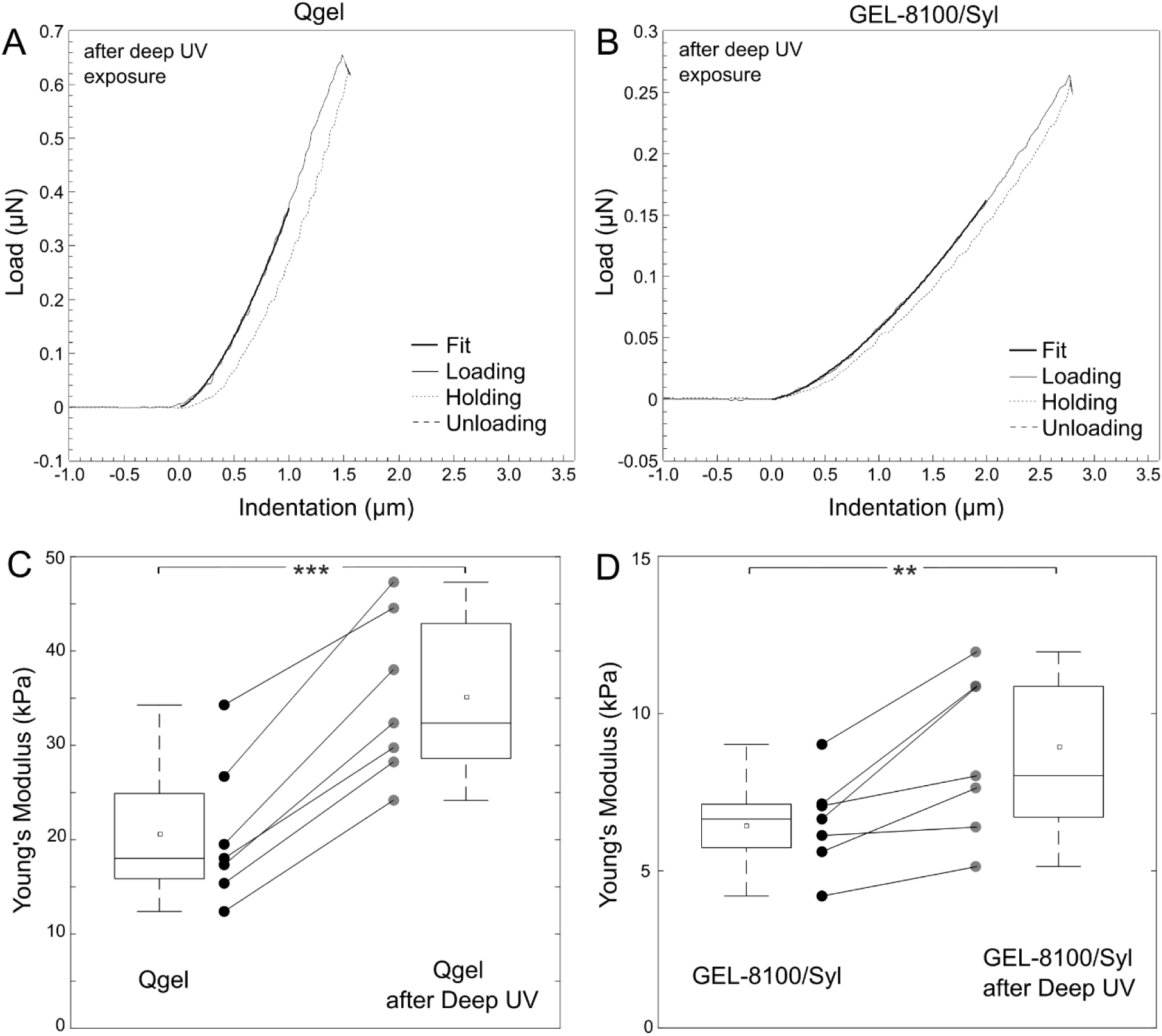
Deep UV exposure stiffens both kinds of soft silicone. (A, B) Load-indentation curves for the indentation of two kinds of soft silicone after deep UV exposure, measured in 1% (w/w) sodium dodecyl sulfate (SDS) in PBS: (A) 1:2.2 Qgel and (B) 1:1 GEL-8100/0.5% Sylgard crosslinking agent. Loading curves are shown as solid lines, unloading curves as dashed lines and the Hertz model fit as bold solid line. (C, D) Box plots of the Young’s moduli of two kinds of soft silicone: (C) 1:1 Qgel and (D) 1:1 GEL-8100/0.5% Sylgard crosslinking agent, measured before and after sample exposure to deep UV. The horizontal line inside the box represents the median, small square inside the box marks the mean, the top and bottom edges of the box represent the 25th and 75th percentile, and the whiskers extend to the most extreme data points. Accompanying individual data from independent samples show deep UV-induced stiffening in each sample. A paired t-test was considered suitable for the data in (C, D) after a Shapiro-Wilk test for normality yielded p values in the 0.30-0.92 range meaning that the null hypothesis that the data are from a normal distribution could not be rejected. The paired t-test p value for (C) was < 0.001 and for (D) was < 0.01, indicating statistically significant differences.

**Figure 4.**
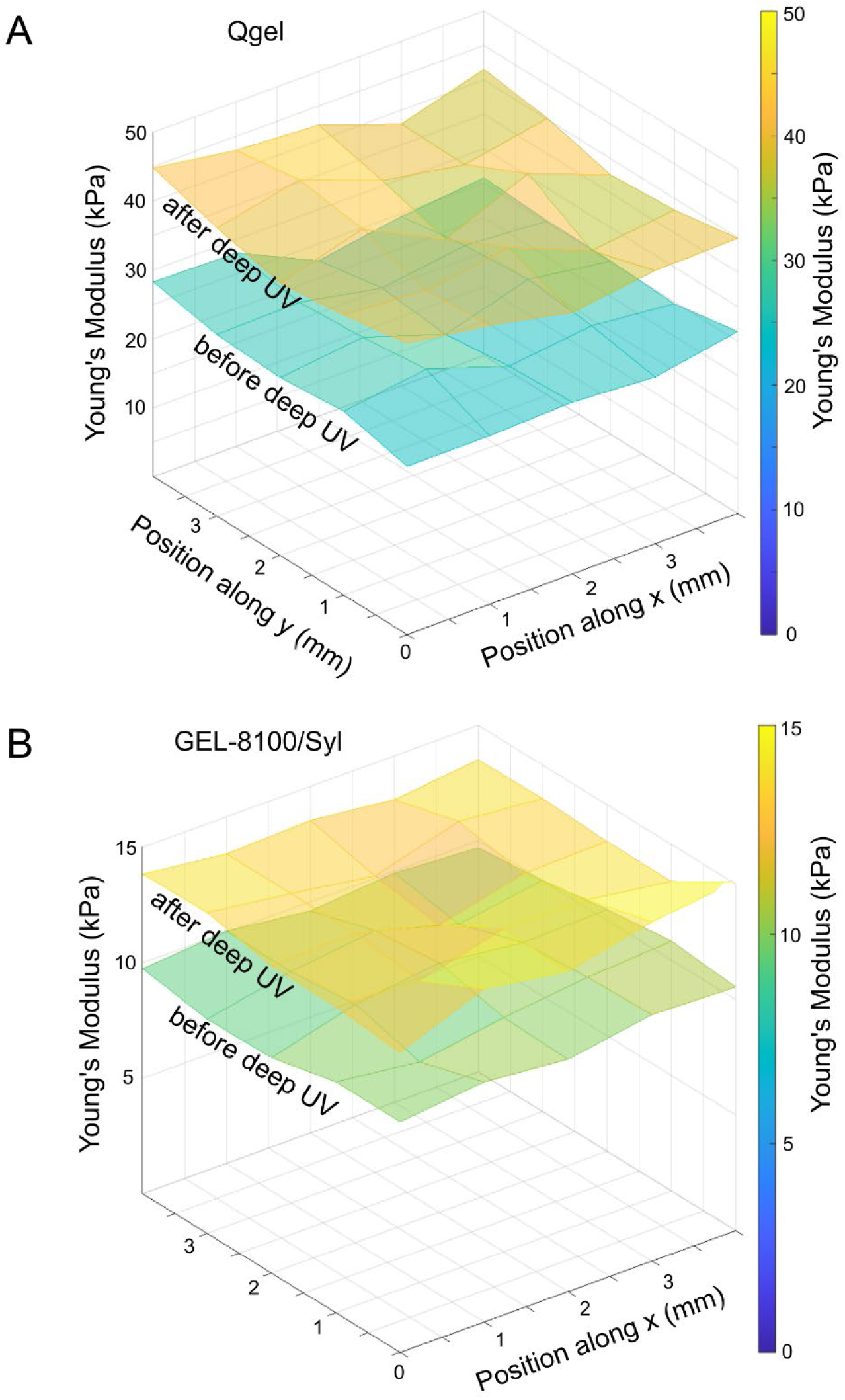
Nanoindentation scans along soft silicone surfaces without and after deep UV exposure. Matrix scans of (A) 1:2.2 Qgel and (B) 1:1 GEL-8100/0.5% Sylgard crosslinking agent, before and after deep UV exposure.

We then wanted to test if the bulk properties of the soft silicones were affected or if the stiffening effect was restricted to the surface as suggested previously with stiffer silicones (30, 31). We therefore used bulk shear rheology to assess whether the bulk mechanical properties of either kind of soft silicone was different after deep UV exposure. Measurement of the shear storage and loss moduli over two decades of angular frequency showed that the bulk mechanical properties were unaffected after deep UV exposure (Fig 5). Prior works with stiffer PDMS (31, 32, 34) have used techniques such as X-ray photoelectron spectroscopy (XPS) (31, 34) and attenuated total reflectance Fourier transform infrared spectroscopy (ATR-FTIR) (32) to probe the surface of deep UV exposed stiff PDMS. XPS in particular revealed (34) that the concentration of elemental oxygen (O) increased relative to carbon (C) and Silicone (Si) upon deep UV exposure of stiff silicone. ATR-FTIR was used to show (32) that a decrease in signal due to -CH_3_ and an increase in signal due to -OH, suggesting oxidation upon deep UV exposure of stiff silicone. We therefore hypothesized that, if the surface of soft silicones were getting oxidized, the mole fraction of oxygen would increase and carbon would decrease upon deep UV exposure. SEM of untreated or deep treated soft silicone surfaces did not show any obvious surface topographical features (Fig 6A,C).

**Figure 5.**
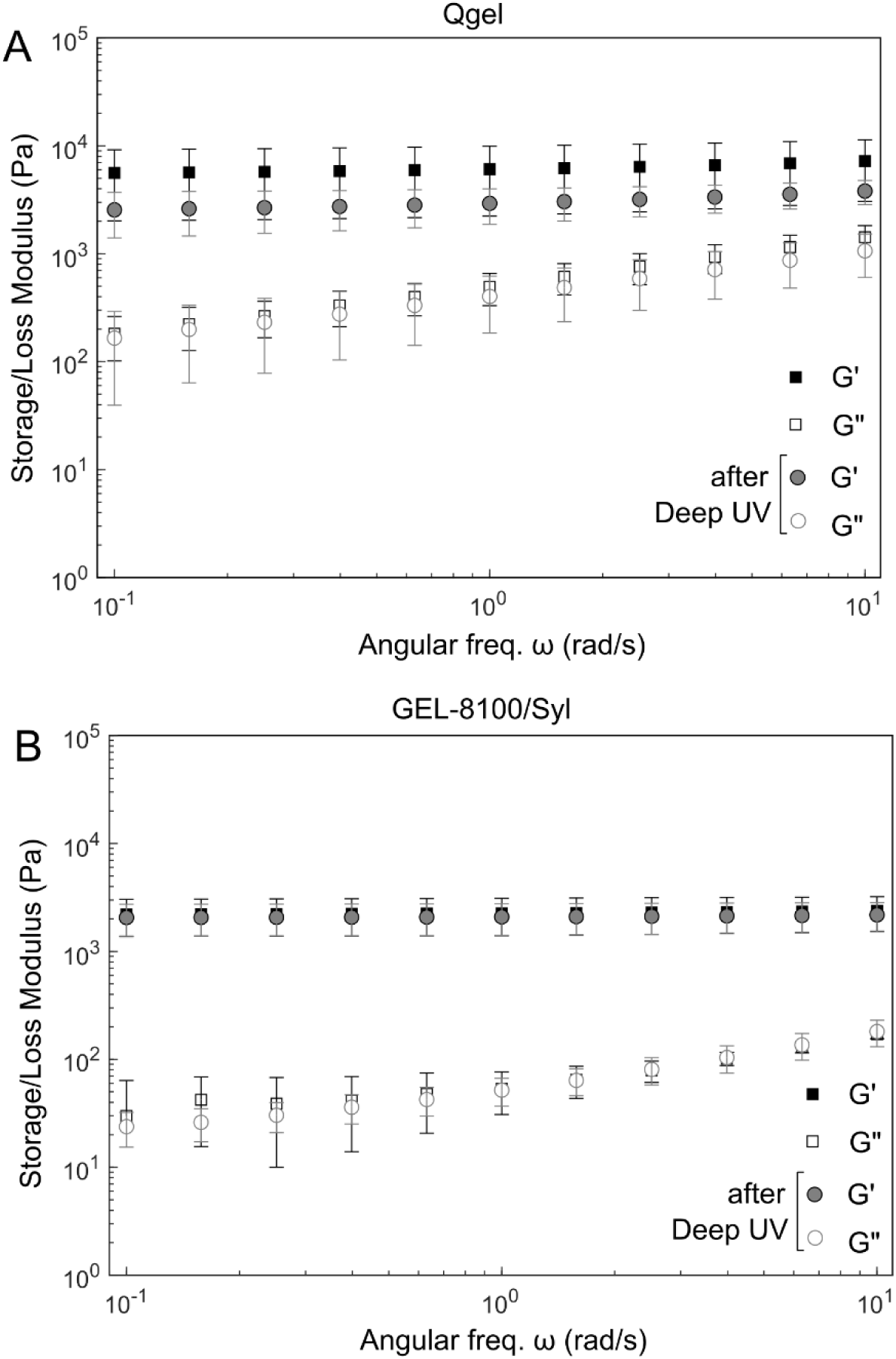
Bulk rheology of soft silicones was unaffected by deep UV exposure. Bulk rheology of two kinds of soft silicone after deep UV exposure: (A) 1:2.2 Qgel and (B) 1:1 GEL-8100/0.5% Sylgard crosslinking agent. Storage shear modulus (G’, filled symbols) and loss shear modulus (G”, open symbols) have been plotted as a function of angular frequency (ω) for either soft silicone without (black symbols) or after deep UV exposure (grey symbols).

**Figure 6.**
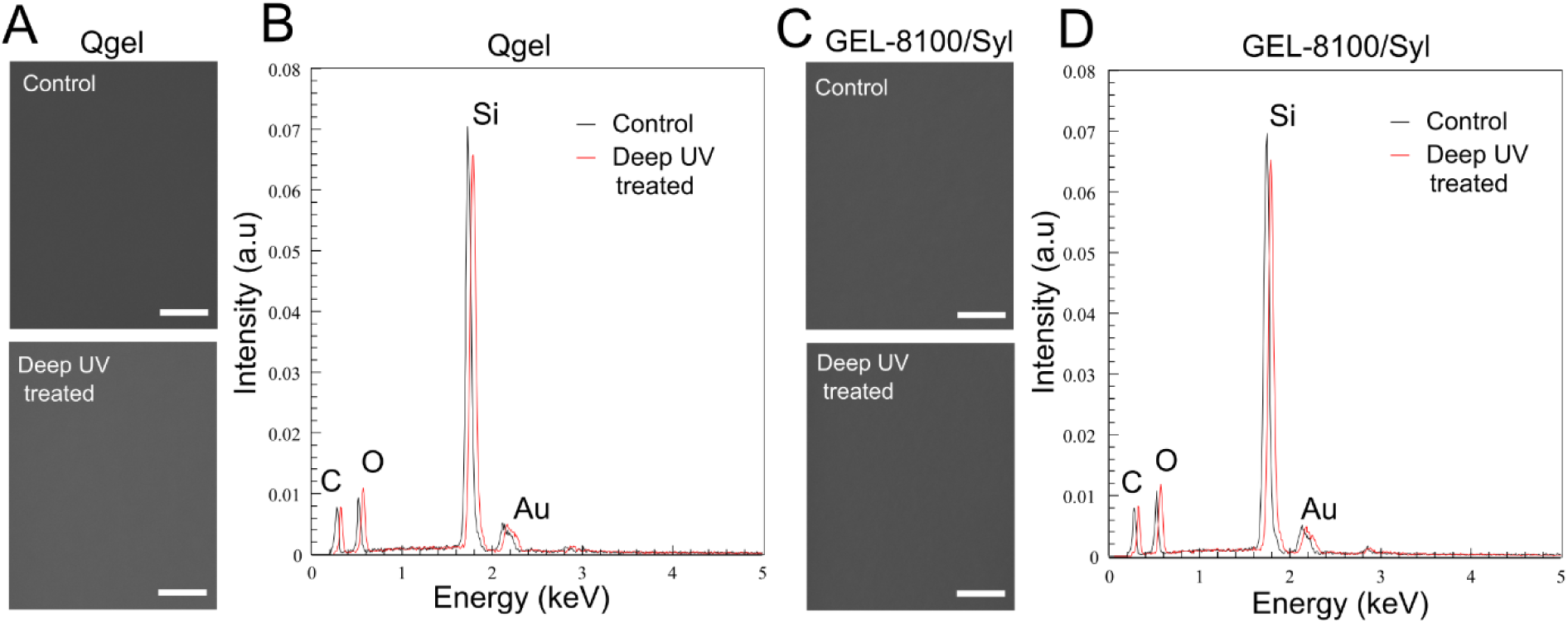
SEM and EDS of soft silicones. SEM and EDS of 1:2.2 Qgel (A, B) and GEL-8100/Syl (C,D). SEM of either kind of soft silicone (A,C) at 30,000x magnification shows a near-featureless smooth surface. Scale bar is 0.5 μm. (B,D) EDS of either kind of soft silicone shows a small but consistent relative change in peak intensities corresponding to an increase in the mole fraction of oxygen and decrease in the mole fraction of carbon. The mole percentages of C, O, Si for untreated Qgel were 37.95 ± 0.81, 25.51 ± 0.86 and 34.77 ± 0.30, for deep UV treated Qgel were 33.03 ± 0.83, 28.93 ± 0.77 and 35.41 ± 0.26, for untreated GEL-8100/Syl were 35.93 ± 0.76, 27.41 ± 0.73 and 34.99 ± 0.25 and for deep UV treated GEL-8100/Syl were 33.36 ± 0.82, 29.61 ± 0.75 and 34.58 ± 0.25. Note that the Au-Pd alloy used to coat the SEM/EDS samples accounted for the rest of the mole percentage, such that the total is 100% when the Au and Pd mole percentages are included. Also note that, in (B,D), the spectrum for deep UV treated case is shifted to the right by 0.05 keV to aid in clarity of comparison with the untreated case. It is the relative change in peak intensity that relates to element abundance.

X-ray spectroscopy (Energy-dispersive X-ray spectroscopy, EDS (38)) (Fig 6B,D) revealed that relative peak intensities changed upon deep UV exposure (similarly for both Qgel and GEL-8100/Syl) such that the mole percentage of oxygen increased from 25.51 ± 0.86 to 28.93 ± 0.77 for Qgel and from 27.41 ± 0.73 to 29.61 ± 0.75 for GEL-8100/Syl. Concomitantly, the mole percentage of carbon decreased from 37.95 ± 0.81 to 33.03 ± 0.83 for Qgel and from 35.93 ± 0.76 to 33.36 ± 0.82 for GEL-8100/Syl upon deep UV exposure, with the rest of the mole percentage corresponding to silicon as well as the gold-palladium coating used in SEM/EDS. Our EDS results are consistent with surface oxidation yielding a layer of silicone oxide of thickness considerably less than the EDS interrogation thickness of ∼ 1 μm, given the small but consistent shifts seen in both kinds of soft silicone.

To further investigate the effect of deep UV treatment on the soft silicones, we obtained the ATR-FTIR spectra of the sample surfaces without and after deep UV treatment. Since the ATR-FTIR signal strength can vary due to the variations in the extent of sample contact, we first normalized the absorbance spectra as detailed in the Materials and Methods section. Both untreated soft silicones display peaks characteristic of PDMS (32) (Fig. 7 A,B): Si–CH_3_ rocking and Si–C stretching at 790-800 cm^-1^ (a), asymmetric Si–O–Si stretching in siloxane doublet at 1000-1100 cm^-1^ (b), symmetric Si–CH_3_ deformations at ∼1260 cm^-1^ (c), and asymmetric Si–CH_3_ stretching at ∼2960 cm^-1^ (d). Surface oxidation is expected to break Si-C and C-H bonds, and lead to the initial formation of Si-OH bonds and then Si-O-Si bonds as in silica or silicon sub-oxide (i.e., SiO_x_ where x < 2). The initial formation of Si-OH bonds should show as a shallow, broad band in 3050–3700 cm^−1^ due to stretching of -OH in Si–OH and more advanced oxidation due to formation of a silica-like layer or silicon sub-oxide layer should show as shifts from a double peak at 1000-1100 cm-1 to more of a broader single peak.

**Figure 7.**
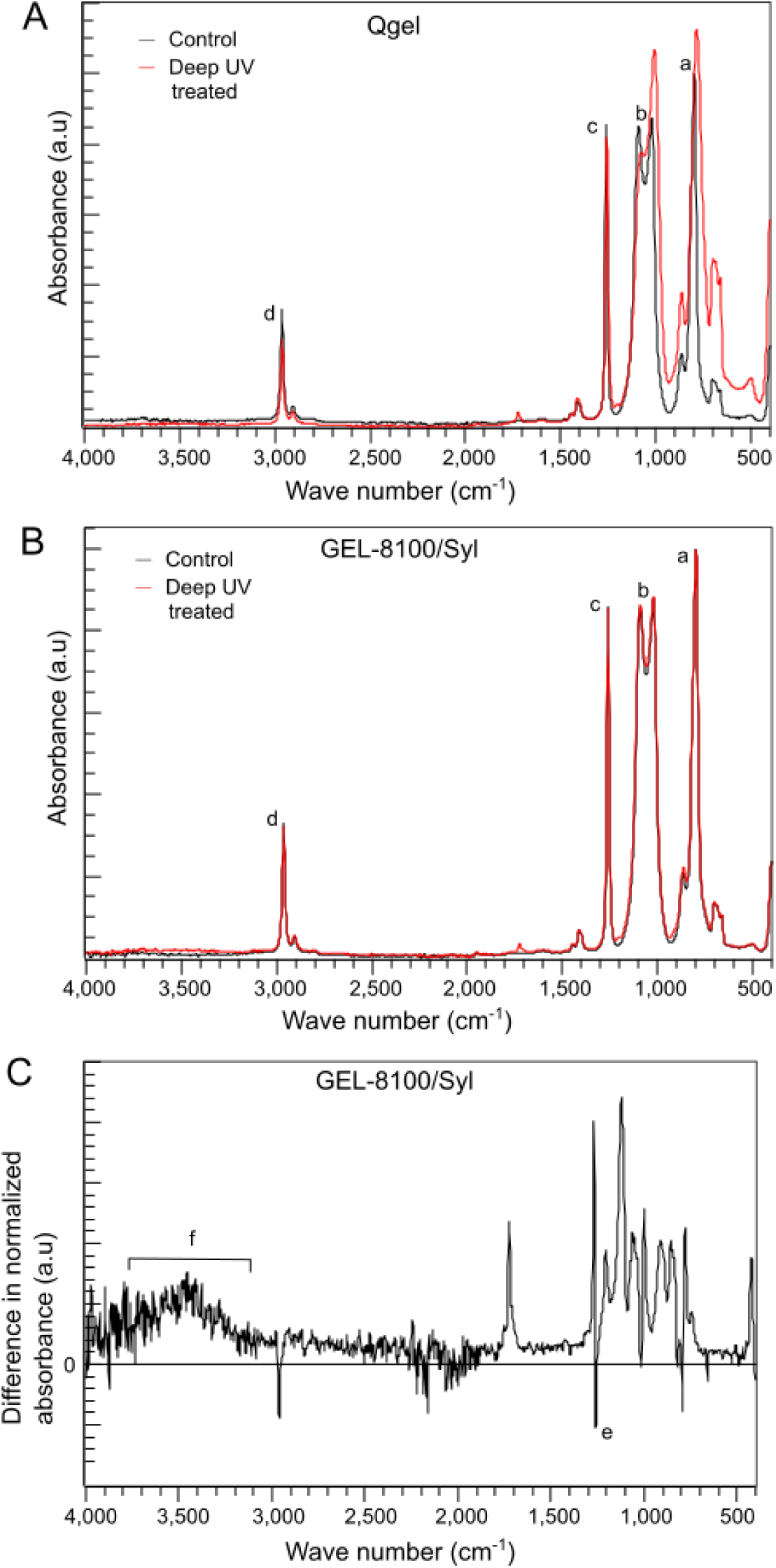
ATR-FTIR spectroscopic characterization of soft silicones treated without or after deep UV exposure. (A) ATR-FTIR absorption spectra of 1:2.2 Qgel without (control, black) or after deep UV treatment (red). Prominent peaks in (A, B) are Si–CH_3_ rocking and Si–C stretching at 790-800 cm^-1^ (a), asymmetric Si–O–Si stretching in siloxane doublet at 1000-1100 cm^-1^ (b), symmetric Si–CH_3_ deformations at ∼1260 cm^-1^ (c), and asymmetric Si–CH_3_ stretching at ∼2960 cm^-1^ (d). Major changes to the asymmetric Si–O–Si stretching siloxane doublet at 1000-1100 cm^-1^ (b) are apparent after deep UV treatment of Qgel. (B) ATR-FTIR absorption spectra of 1:1 GEL-8100/0.5% Sylgard crosslinking agent without (control, black) or after deep UV treatment (red). Changes upon deep UV treatment are less apparent here, and therefore the difference in spectra between deep UV treated and untreated GEL-8100/Syl is presented in (C). (C) Difference between the normalized absorbance spectra of deep UV treated GEL-8100/Syl and untreated GEL-8100/Syl. Broad increase in signal at 3050–3700 cm^-1^ due to stretching of -OH in Si–OH and a decrease in the signal due to symmetric Si–CH_3_ deformations at 1260 cm^-1^ are now apparent.

The difference in ATR-FTIR spectra is quite apparent for Qgel upon deep UV treatment (Fig. 7A). The main Si-O-Si double peak at 1000-1100 cm^-1^ (b) broadens and shifts to a single peak closer to 1000 cm^-1^ as expected for a silicon sub-oxide layer (42). Symmetric Si–CH3 deformation at ∼1260 cm^-1^ also shows a decrease, as expected due to Si-C bond breakage. The difference in ATR-FTIR spectra is, however, more subtle for GEL-8100/Syl upon deep UV treatment (Fig. 7B). In order to more clearly observe the change in spectra, the difference between the normalized absorbance spectra of deep UV treated GEL-8100/Syl and untreated GEL-8100/Syl has been plotted in Fig. 7C. Few key features are apparent here: The Si−CH3 symmetric deformation signal at 1260 cm^−1^ decreases (marked as e in Fig. 7C) as expected due to oxidation, and a shallow broad increase appears between 3050–3700 cm^−1^ (marked as f in Fig. 7C), corresponding to Si−OH group formation due to oxidation (32). Thus, Qgel shows higher levels of oxidation (due to evidence of silica like silicon sub-oxide layer (42)) compared to GEL-8100/Syl that shows evidence milder oxidation via Si-OH group formation. This results also agrees with the greater stiffening effect seen in Qgel compared to GEL-8100/Syl. To assess if the surfaces have become more hydrophilic upon deep UV treatment, we performed water contact angle measurements on either soft silicone surface without and after deep UV treatment. We found that Qgel’s water contact angle decreased from 110^0^±10^0^ to 98^0^±8^0^ after deep UV treatment (p < 0.01) and the contact angle of GEL-8100/Syl decreased from 99^0^±3^0^ to 92^0^±2^0^ after deep UV treatment (p < 0.05). Thus, contact angle measurements show that the surfaces became more hydrophilic after deep UV treatment, as expected due to oxidation.

Given that we determined a change in the apparent Young’s moduli after deep UV exposure, and given that this is a surface effect, we considered that a top layer of thickness *t* had been stiffened to a higher Young’s modulus E_surf_ (by deep UV in air) which is much higher than the bulk Young’s modulus E_bulk_. Thus, we effectively have an elastic bilayer, whose overall apparent Young’s modulus is E_app_. The Hertz indentation of an elastic bilayer has been modelled before (43) and we can thus roughly estimate E_surf_ from E_app_ that we measured. It has been shown (43) that the ratio (E_app_ – E_surf_)/(E_bulk_ – E_surf_) has a sigmoidal dependence on *a*/*t*, where *a* is the contact radius and *t* is the thickness of the top layer that has been stiffened. If we assume that the stiffened region is the top 50 nm (31, 44), then, for our measurements, *a* ∼ 6 µm, *t* ∼ 0.05 µm, so *a*/*t* ∼ 120 and therefore (E_app_ – E_surf_)/(E_bulk_ – E_surf_) is ∼0.99 (43). This leads to estimates of E_surf_ of ∼ 1500 kPa for Qgel and 200 kPa for GEL-8100/Syl. These estimates suggest that the the top surface has actually stiffened by a factor of 30-75, even though the apparent moduli have only increased by tens of percent. One limitation of the deep UV treatment method employed here as it applies to mechanobiology studies is that the actual local Young’s modulus is expected to vary as a function of depth from the surface of the soft silicone, which implies that the length scale of the cell collective will determine the effective Young’s modulus sensed. Change in surface composition upon deep UV exposure also implies that any protein coupling strategies will have to be customized to consider equal protein coverages between untreated and deep UV stiffened substrates. The deep UV exposure conditions will also have to be optimized depending on the application in order to obtain desired levels of substrate stiffness alteration.

## Conclusions

To conclude, we have used nanoindentation to assess the effect of deep UV treatment in the presence of air (in effect, UVO treatment) on the mechanical properties of two kinds of soft silicones (Qgel and GEL-8100/Syl) of particular interest in mechanobiology studies. We first validated the utility of using a surfactant (41) to eliminate adhesion between glass probes and both types of soft silicones. We then found that deep UV treatment reliably increases the apparent Young’s moduli of either kinds of soft silicone by tens of percent. Bulk rheology showed that the change in apparent Young’s moduli is due to a surface effect, in agreement with prior studies with stiffer PDMS silicones (Sylgard 184). SEM of either kind of soft silicone showed no major discernible topographical features, consistent previously reported results with GEL-8100 of a different composition (14). EDS (38) of both kinds of soft silicone showed an increase in the mole fraction of oxygen and decrease in the mole fraction of carbon, consistent with surface oxidation yielding a silicon oxide layer of fraction of a micron thickness. We also used ATR-FTIR to show that, while the level of Si-CH_3_ bonds decreased in both Qgel and GEL-8100/Syl, the extent of oxidation was different in that a silica-like silicon sub-oxide layer formed in Qgel whereas Si-OH groups were formed in GEL-8100/Syl. We used an elastic bilayer model to estimate the stiffness of the surface layer and found that its Young’s modulus is several hundred kPa (GEL-8100/Syl) or over 1 MPa (Qgel). Despite being large compared to bulk soft silicone moduli, these values are considerably smaller than the reported moduli of stiff layers formed upon UVO treatment of Sylgard 184 based stiffer PDMS (34).

Cell-sensing of matrix stiffness is of fundamental importance in normal physiology and pathology (45–49). Since soft silicones coated with matrix molecules have found extensive use as substrates for cell culture in mechanobiology studies (50), the stiffening effect reported here upon deep UV treatment in the presence of air can be used to inform their applications in mechanobiology studies in several ways: (i) Deep UV exposure of soft silicone substrates can be spatially controlled with the use of synthetic quartz masks wherein a spatially varying chromium coating specifies where deep UV can pass through and where it cannot (51). The creation of surfaces with spatially micropatterned effective Young’s moduli, by spatially controlled deep UV exposure can enable the study of the effect of the spatial variation of substrate stiffness on cell migration and multi-cellular spatial organization. (ii) Cells like chondrocytes (52) are adapted to relatively higher matrix/substrate stiffness. The mechanosensing behaviors of such cells, whose physiologically relevant tissue moduli match the surface stiffness of deep UV exposed soft silicone, can be studied using soft silicone substrates to which matrix molecules can be coupled after deep UV treatment (36) (iii) The mechanosensing behaviors of supra-cellular assemblies like large epithelial islands/monolayers, that can sense beyond the stiffened layer of deep UV exposed soft silicone can also be studied with these substrates, provided their physiologically relevant matrix moduli match the overall apparent Young’s moduli of deep UV exposed soft silicone (48) which we have shown here to be 30-70% higher than that of untreated soft silicones. (iv) Finally, cell-cell contacts are also mechanosensitive interfaces (53). Since deep UV treatment stiffens soft silicones as shown here, biomimetic approaches to mimic cell-cell contact interfaces (14) can potentially use this treatment to develop interfaces whose surface stiffening can mimic cell-cell junction regions that have stiffened due to actomyosin enrichment at the contact. Alternately, our study also informs that one should avoid using deep UV to chemically modify soft silicone surfaces to couple biomolecules (36), when any stiffening effect is undesirable. In conclusion, we have thus demonstrated a simple physical method to alter the stiffness of soft silicones and propose that this approach can be widely inform mechanobiological studies.

## Acknowledgments

We thank Wei Cao at the ODU Applied Research Center at Newport News, VA for assistance with SEM and EDS. We thank Prof. Bala Ramjee for assistance with ATR-FTIR.

## Author contributions

V.M. designed research; A.W., Z. B., C.O., S.S., and V.M. performed research; A.W. and V.M. wrote the paper.

